# Fendioxypyracil, a new and systemic PPO-inhibiting herbicide for X-spectrum weed control

**DOI:** 10.64898/2026.01.05.697680

**Authors:** Tobias Seiser, Aimone Porri, Philipp Johnen, Silke Zeyer, Judith Wahrheit, Michael Betz, Breght Vandenberghe, Scott Asher, Liliana Parra Rapado

**Affiliations:** BASF SE, Speyererstr, 2, 67117 Limburgerhof, Germany; BASF Corporation, 26 Davis Drive, Research Triangle Park, 27709-3528 United States; BASF Belgium, Technologiepark 101, Gent Oost-Vlaanderen (VLG) 9052 Belgium

## Abstract

**Background:** Fendioxypyracil is a novel protoporphyrinogen oxidase (PPO)–inhibiting herbicide (HRAC Group 14) developed to address the increasing prevalence of herbicide-resistant weeds and to expand available weed control options. PPO inhibitors disrupt chlorophyll biosynthesis by blocking the conversion of protoporphyrinogen IX to protoporphyrin IX, resulting in light-promoted formation of reactive oxygen species and rapid plant necrosis. Building on established PPO chemistry, fendioxypyracil incorporates a pyridine-based core and an aryloxy sidechain designed to enhance binding affinity and post-emergence activity.

**Results:** Greenhouse evaluations demonstrated high efficacy of fendioxypyracil against key grass weeds, including wild oat (*Avena fatua*), crabrass (*Digitaria Sanginalis*), goosegrass (*Eleusina indica*) and barnyard grass (*Echinochloa crus-galli*), as well as strong control of major broadleaf species. Physiological characterization and enzyme inhibition assays confirmed PPO as the primary site and mode of action, with IC₅₀ values lower than those of the commercial standard saflufenacil for both PPO isoforms.

**Conclusion:** Fendioxypyracil represents a next-generation PPO inhibitor with broad-spectrum and systemic activity, offering a valuable new tool for integrated weed management. Its high biological activity and efficacy across multiple weed taxa supports its potential to enhance on-farm weed control strategies and contributes to resistance management programs.

## 1. Introduction

Protoporphyrinogen oxidase (PPO) inhibitor herbicides – classified as Group 14 by the Herbicide Resistance Committee (HRAC) – play a crucial role in modern weed management, providing effective control of broadleaf and some grass weeds across diverse cropping systems. These herbicides are valued for their rapid action, often producing visible symptoms within hours under favorable environmental conditions (1). Their versatility allows for both pre-and post-emergence applications, either as stand-alone products or in tank mixtures, supporting both efficacy and resistance management strategies (2).

PPO inhibitors act by targeting two key enzymes in the tetrapyrrole biosynthetic pathway, PPO 1 and 2, which are involved in the synthesis of chlorophyll and heme – two compounds vital for plant survival. Inhibition of these enzymes disrupts the conversion of protoporphyrinogen IX to protoporphyrin IX, leading to the accumulation of toxic intermediates. Under light and aerobic conditions, this results in the strong generation of reactive oxygen species (ROS), causing rapid membrane damage, tissue necrosis, and ultimately plant death (3).

Several chemical families fall under PPO inhibitors, including diphenylethers (e.g., fomesafen), N-phenyltriazolinones (e.g., sulfentrazone), and N-phenylimides (e.g., saflufenacil (4)). Despite their structural diversity, these herbicides share a common mode of action (MoA) and are widely used in crops such as soybean, corn, cotton, and cereals. Their high efficacy, rapid symptom development, and relatively short environmental persistence make them attractive for both conventional and conservation tillage systems.

Continuous innovation in PPO inhibiting chemistry has led to the discovery of new active ingredients, even decades after the first introduction of nitrofen in 1964. Recent advances include the development of molecules with improved characteristics, such as tiafenacil (launched in 2020) (5). However, the emergence of herbicide resistance and the need for broader-spectrum solutions drive ongoing research in this area (6, 7).

In response to these challenges, we introduce fendioxypyracil – provisionally approved by ISO - a novel PPO inhibitor designed for broad-spectrum, post-emergence weed control in major crops. Here, we present details on the discovery and synthesis, MoA confirmation, and greenhouse efficacy of fendioxypyracil, highlighting its value as next-generation tool for sustainable weed management (8).

PPO herbicides generally provide strong dicot control both pre-and post-emergence (9). For example, saflufenacil (Kixor®) is an excellent tool for post-emergence conyza control, one of the most difficult dicot weeds to manage (10). Trifludimoxazin (Tirexor®) proved to be exceptionally effective in controlling even PPO-resistant amaranth weeds (11). With the horizon of PPO-tolerant crops in the midterm future and expected, as well as already observed, resistance issues for Glyphosate on grass weeds (12, 13), we aimed to develop a new post-emergence, broad-spectrum PPO herbicide – a crucial tool for future weed management.

In the late 1990s, Novartis (now Syngenta) published PPO inhibitor herbicides bearing pyridinone (Scheme 1, **1**) as well as pyridine cores (**2**) (14–16). The pyridine motif was further explored by the newly formed Syngenta company in the early 2000s, resulting in pyrido-oxazinone (Scheme 1, **3**) and 3-arylpyridine structures (**4**) (17–19). None of these structures were commercialized, and the central pyridine motif in PPO structures did not receive further attention until BASF revisited it in 2017 (8). By employing nucleophilic aromatic substitution (SNAr) reactions, diverse nucleophiles can be readily introduced onto the pyridine core. Even complete warheads bearing a nucleophilic nitrogen can be directly attached (20). Besides warheads, sidechains connected via nucleophilic atoms like oxygen can be easily introduced. For PPO herbicides, ether sidechains proved to be highly active, ranging from simple methoxy-or propargyloxy-ethers to lactic acid-and acetal-sidechains (9, 21). Another interesting class of ether sidechains includes substituted aryloxy residues. In the early 2000s, Sumitomo described a set of phenoxy-, pyridyloxy-, and pyrimidyloxy-sidechains with diverse substitutions in the ortho-, meta-, or para-positions (22, 23).

Sumitomo’s work on aryloxy sidechains ultimately led to the PPO market product Epyrifenacil (Rapidicil®) (24), which features a pyridyl-ether sidechain and a phenyl core. Connected through SNAr reactions, the aryloxy side chains are particularly intriguing when combined with the pyridine core. The preferred dihedral angle between the PPO core and the aryloxy side chain is more constrained with a pyridine core compared to a phenyl core (Figure 1). This pyridine core induced preorganization matches the ligand’s dihedral angle in the enzyme pocket more closely, thereby promoting a favorable fit with the targeted binding-mode geometry. While, in general, ortho-, meta-, and para-substituted phenoxy as well as pyridyloxy residues show good biological activity in combination with the pyridine core (19), a simple catechol unit elongated by ethyl acetate provided excellent grass and dicot control post-emergence. This led to the invention of compound **5**, provisionally approved by ISO as fendioxypyracil, BASF’s next-generation herbicide for post-emergence broad-spectrum weed control (Scheme 1, **5**). Here, we present evidence for the high efficacy of fendoxypyracil and the mode and site of action of fendioxypyracil.

**Figure 1:**
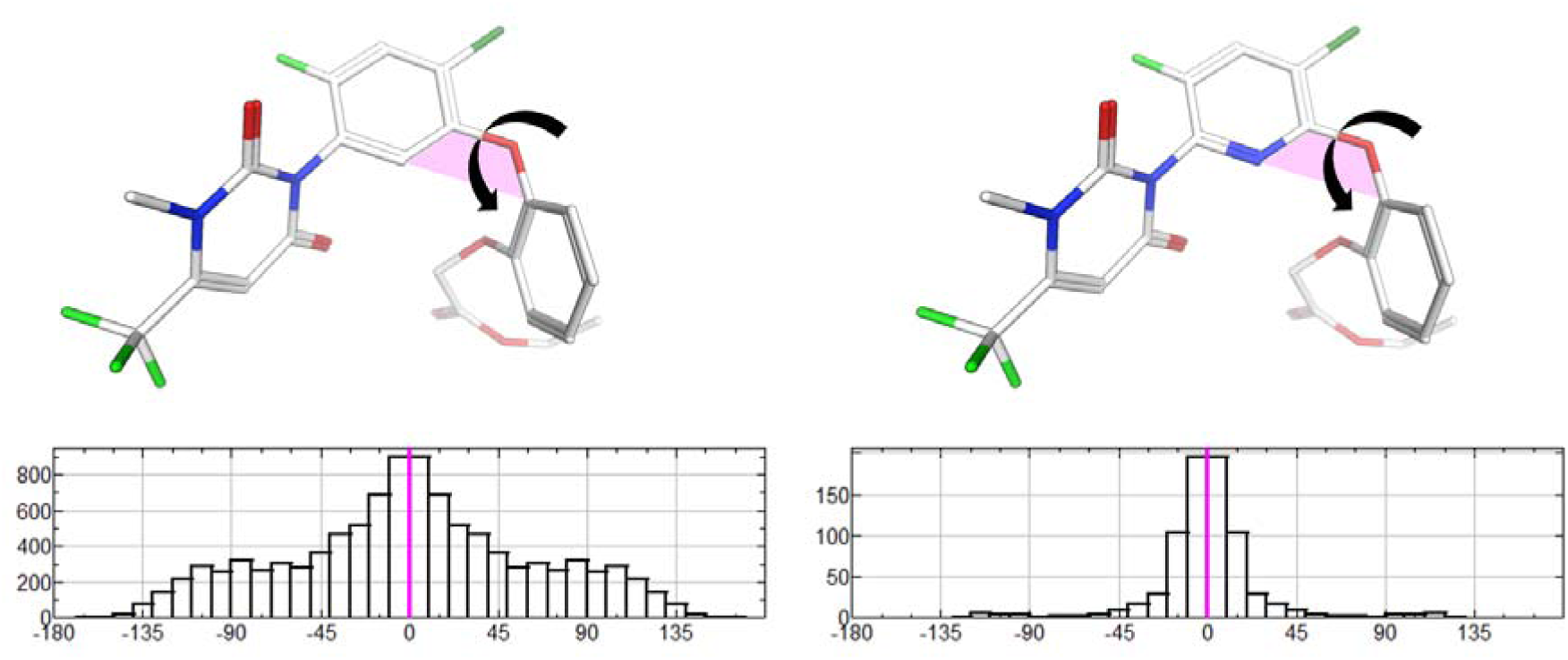
Histograms of the dihedral angle distributions for the phenyl-core (left) and pyridine-core (right) variants. The 3D structures, which correspond to the conformations observed in the protein binding pocket, are shown in stick representation. Carbon atoms are colored white, nitrogen blue, oxygen red, chlorine dark green, and fluorine light green. The angles shown in the histograms are indicated in the structures by black arrows. 0° (magenta line in the histograms) means the C–C–O–C (phenyl) or N–C–O–C (pyridine) atoms lie in the same plane (shown as magenta plane in the 3D structures).

## 2. Material and Methods

### 2.1 Statistical distribution of dihedral angle in Figure 1

The histogram of the statistical distribution of dihedral angles was generated with the software MOE (25).

The data were obtained from the Mogul program (CSD 6.00 with CSD Aug25 update) of the CCDC suite (26).

The analyzed dihedral angles were C–C–O–C for the phenyl variant and N–C–O–C for the pyridine variant.

### 2.2 Synthetic Method

#### 2.2.1 Synthesis of Fendioxypyracil (**5**)

##### Step (a): 2-azido-6-(2-benzyloxyphenoxy)-5-chloro-3-fluoro-pyridine (**7**)

To a solution of 5.0 g (29 mmol) 3-chloro-2,5,6-trifluoropyridine (CAS 2879-42-7) in 50 mL DMSO was added 2.1 g (33 mmol) NaN_3_ and the solution was stirred at room temperature for 3 hours. Then 19.5 g (60 mmol) Cs_2_CO_3_ was added followed by a solution of 6.2 g (31 mmol) 2-(Benzyloxy)phenol) in 40 mL DMSO. The mixture was stirred at room temperature for 16 hours, water (200 mL) was added and the mixture was extracted with ethyl acetate (3*300 mL). The organic layer was separated, washed with brine, dried over anhydrous Na_2_SO_4_, filtered and the solvent was removed under reduced pressure. The crude material (15 g) was used without further purifiction in the next step. [M+H] = 371.0; Rt = 1.368 min

##### Step (b): 2-amino-6-(2-benzyloxyphenoxy)-5-chloro-3-fluoro-pyridine

To a solution of 15 g of compound **7** in THF (100 mL) was added 9.7 g (150 mmol) zinc and 100 mL aq. NH_4_Cl dropwised at 0°C. The mixture was stirred for 16 hours at room temperature, filtered and the filter cake was washed with ethyl acetate (50 mL). The filtrate was extracted with ethyl acetate (3*200 mL), the combined organic layer was dried over anhydrous Na_2_SO_4_, filtered and the solvent was removed under reduced pressure. The crude material was purified by silica gel column (petrol ether/ethyl acetate) to give 8.8 g (25.6 mmol, 88% from 3-chloro-2,5,6-trifluoropyridine) of 2-amino-6-(2-benzyloxyphenoxy)-5-chloro-3-fluoro-pyridine. [M+H] = 345.0; Rt = 1.232 min

##### Step (c): ethyl N-[6-(2-benzyloxyphenoxy)-5-chloro-3-fluoro-2-pyridyl]carbamate (**8**)

To a solution of 8.8 g (25.6 mmol) of 2-amino-6-(2-benzyloxyphenoxy)-5-chloro-3-fluoro-pyridine in 80 ml dichloromethane was added 3 g (38 mmol) pyridine followed by 4 g (37.5 mmol) ethyl chloroformate. The mixture was stirred at 25°C for 20 hours, diluted with water and extracted with dichloromethane. The combined organic layer was washed with brine, dried over anhydrous Na_2_SO_4_ and concentrated to give 14.4 g of a mixture of carbamate **8** and the di-substituted derivative. The crude mixture (12.4 g) was dissolved in 200 ml ethanol and aqueous NaOH (1 M) was added dropwise at 0°C with stirring. The mixture was stirred at 15°C for 6 hours, diluted with brine and extracted with ethyl acetate. The combined organic layer was dried over anhydrous Na_2_SO_4_, filtered and the solvent was removed under reduced pressure. The crude product was purified by column chromatography on silica (petrol ether/ethyl acetate) to give 6.6 g (15.9 mmol, 62%) of the desired compound **8**. [M+H] = 417.1; Rt = 1.293 min

##### Step (d): 3-[6-(2-benzyloxyphenoxy)-5-chloro-3-fluoro-2-pyridyl]-6-(trifluoromethyl)-1H-pyrimidine-2,.4-dione

To a solution of 1.7 g (43 mmol) NaH in NMP (60 mL) at 0°C was added 6 g (14 mmol) of compound **8** and the mixture was stirred for 30 minutes at 35°C. Then 3.9 g (21 mmol) of ethyl (E)-3-amino-4,4,4-trifluoro-but-2-enoate (CAS: 372-29-2) was added and the reaction mixture was stirred at 100°C for 3 days. The resulting mixture was quenched with ice water (100mL), acidified to pH=2 by using 6N HCl and extracted with ethyl acetate (3*100mL). The combined organic layer was washed with brine, dried over anhydrous Na_2_SO_4_, concentrated and directly used in the next step. [M+H] = 508.0; Rt = 1.240 min

##### Step (e): 3-[6-(2-benzyloxyphenoxy)-5-chloro-3-fluoro-2-pyridyl]-1-methyl-6-(trifluoromethyl)-pyrimidine-2,4-dione (**9**)

To a solution of 6.5 g (12.8 mmol) of 3-[6-(2-benzyloxyphenoxy)-5-chloro-3-fluoro-2-pyridyl]-1-methyl-6-(trifluoromethyl)-pyrimidine-2,.4-dione in 65 mL acetonitrile was added 5.3 g (38 mmol) K_2_CO_3_ followed by 7.3 g (51 mmol) methy iodide at 0 ° C with stirring. The mixture was stirred at 15^°^C for 16 hours, then water (80 mL) was added, and the pH was adjusted to pH=5 by using 2M HCl. The mixture was extracted with ethyl acetate (3*90mL), the combined organic layer was washed with brine and dried over anhydrous Na_2_SO_4_, filtered and the solvent was removed under reduced pressure yielding 7 g of the crude product **9**, which was used without further purification.

^1^H-NMR (CDCl_3_, ppm): 7.63 (d, J=7.28 Hz, 1 H); 7.21 - 7.25 (m, 4 H); 7.12 - 7.17 (m, 2 H); 6.98 (t, J=7.03 Hz, 3 H); 6.26 (s, 1 H); 4.99 (s, 2 H); 3.47 (s, 3 H). [M+H] = 522.0; Rt = 1.323min

##### Step (f): 3-[5-chloro-3-fluoro-6-(2-hydroxyphenoxy)-2-pyridyl]-1-methyl-6-(trifluoromethyl) pyrimidine-2,4-dione

To a solution of 7 g (13.4 mmol) of compound **9** in 70 mL xylene was added 3.6 g (26 mmol) solid AlCl_3_ at 15°C with stirring. The mixture was stirred at 130 ° C for 16 hours and after cooling to 15°C, ice-water (100 mL) was added to the mixture. After separation of the xylene layer, the water phase was extracted with ethyl acetate (3*80 mL), the combined organic layer was dried over anhydrous Na_2_SO_4_, filtered and the solvent was removed under reduced pressure. The crude product was purified by column chromatography on silica gel (petrol ether/ethyl acetate) to give 3.2g (7.4 mmol, 55%) of 3-[5-chloro-3-fluoro-6-(2-hydroxyphenoxy)-2-pyridyl]-1-methyl-6-(trifluoromethyl)-pyrimidine-2,4-dione.

^1^H-NMR (CDCl_3_, ppm): 7.80 (d, J=7.26 Hz, 1H); 7.03 – 7.19 (m, 3H); 6.93 (dt, J=7.68 Hz, J=1.7 Hz, 1H); 6.3 (s, 1H); 5.6 (s, 1H); 3.5 (s, 3H).[M+H] = 431.9; Rt = 1.077 min

##### Step (g): ethyl 2-[2-[[3-chloro-5-fluoro-6-[3-methyl-2,6-dioxo-4-(trifluoromethyl)pyrimidin-1-yl]-2-pyridyl]oxy]phenoxy]acetate, Fendioxypyracil, (**5**)

To a solution of 0.2 g (0.46 mmol) of 3-[5-chloro-3-fluoro-6-(2-hydroxyphenoxy)-2-pyridyl]-1-methyl-6-(trifluoromethyl)-pyrimidine-2,4-dione in 10 mL dry acetonitrile was added 0.19 g (1.3 mmol) K_2_CO_3_ at 0°C followed by dropwise addition of 0.15 g (0.92 mmol) ethyl bromoacetate. The mixture was stirred at 15°C for 16 hours, diluted with 15 ml water and extracted with ethyl acetate (3*15mL). The combined organic layer was washed with brine, dried over anhydrous Na_2_SO_4_, filtered and the solvent was removed under reduced pressure. The crude product was purified by reversed phase preparative HPLC containing TFA to give 0.16 g (0.31 mmol, 67%) of ethyl 2-[2-[[3-chloro-5-fluoro-6-[3-methyl-2,6-dioxo-4-(trifluoromethyl)pyrimidin-1-yl]-2-pyridyl]oxy]phenoxy]acetate (**5**, Fendioxypyracil).

^1^H-NMR (CDCl_3_, ppm): 7.76 (d, J=7.28 Hz, 1 H); 7.22 (d, J=7.72 Hz, 1 H); 7.17 (t, J=7.83 Hz, 1 H); 6.99 - 7.06 (m, 1 H); 6.88 (d, J=7.94 Hz, 1 H); 6.25 (s, 1 H); 4.49 (s, 2 H); 4.19 (q, J=7.20 Hz, 2 H); 3.47 (s, 3 H); 1.25 (t, J=7.17 Hz, 3 H). [M+H] = 518.0; Rt = 1.217 min

### 2.3 Biochemistry test Methods

#### 2.3.1 Recombinant Expression, Purification, and In Vitro Inhibition Assays of wild type

##### Amaranthus tuberculatus PPO1 and PPO2 enzymes

The complete coding sequences of wild-type *Amaranthus tuberculatus* PPO1 and PPO2 were synthesized de novo and inserted into the pRSetB expression vector (Invitrogen, Carlsbad, CA, USA) using BamHI and HindIII restriction sites. To facilitate purification, an N-terminal hexahistidine tag was included. Recombinant constructs were transformed into *Escherichia coli* strain BL21(DE3)pLysS (Novagen, EMD Millipore, Billerica, MA, USA), and transformants were selected on LB agar plates containing 100 µg mL⁻¹ ampicillin and 34 µg mL⁻¹ chloramphenicol.

For protein expression, a single colony was inoculated into 3 mL LB medium with antibiotics and incubated at 37 °C with shaking (200 rpm) for 6 h. A 20-µL aliquot of this starter culture was transferred into 20 mL fresh LB medium and grown overnight. The following day, 100 µL of the overnight culture was inoculated into 100 mL ZYM-5052 autoinduction medium supplemented with antibiotics. Cultures were incubated at 37 °C for 5 h and then shifted to 25 °C for an additional 21 h.

Cells were collected by centrifugation at 6,000 × g for 30 min at 4 °C. Pellets were resuspended in PPO lysis buffer [10 mL g⁻¹ pellet; 50 mM NaH₂PO₄, 100 mM NaCl, 5 mM imidazole, 5% (v/v) glycerol, pH 7.5] supplemented with 20 mg mL⁻¹ lysozyme, 30 U mL⁻¹ DNase I, and protease inhibitors (complete EDTA-free, Roche Diagnostics, Mannheim, Germany). Suspensions were sonicated on ice (3 min total, 30 s bursts at 90% amplitude). After centrifugation at 38,000 × g for 30 min at 4 °C, the supernatant was collected and supplemented with 2 mL of 5 M NaCl.

For affinity purification, a 500-µL bed volume of HisPur Ni-NTA resin (Thermo Fisher Scientific, IL, USA) was equilibrated with buffer (20 mM NaH₂PO₄, 50 mM NaCl, 5 mM imidazole, 5 mM MgCl₂, 17% glycerol, 0.1 mM EDTA, pH 8.0). The clarified extract was applied to the resin, which was then washed with 5.6 mL wash buffer (20 mM NaH₂PO₄, 50 mM NaCl, 5 mM imidazole, 17% glycerol, pH 7.5). Bound proteins were eluted with 1 mL elution buffer (20 mM NaH₂PO₄, 50 mM NaCl, 250 mM imidazole, 17% glycerol, pH 7.5). Protein concentrations were measured using a Scandrop nano volume spectrophotometer (Analytikjena, Life Science, Germany). Purity and solubility were confirmed by SDS–PAGE (10%) with 2.5 µg protein per lane.

Enzyme activity assays for PPO1 and PPO2 were performed using a fluorescence-based approach (excitation 405 nm, emission 630 nm). Reactions (187 µL) contained 100 mM Tris–HCl, 1 mM EDTA, 5 mM DTT, 0.0085% Tween 80, and 15 µL enzyme in resuspension buffer (50 mM Tris–HCl, pH 7.3, 3.2 mM EDTA, 20% v/v glycerol). Enzyme concentrations were adjusted for each variant to normalize maximum fluorescence in the absence of inhibitor under saturating substrate conditions.

Dose–response assays were carried out with fendioxypyracil and saflufenacil, dissolved in 80% DMSO. Ten concentrations (10 µL; 1 × 10⁻ to 5.12 × 10⁻¹² M) were tested in the assay mixture, with a 30-min pre-incubation at room temperature before addition of protoporphyrinogen IX. Fluorescence was monitored for 30.25 min (33 cycles of 55 s each) using a CLARIOstar microplate reader (BMG LabTech, Germany). Percent inhibition was calculated relative to untreated (positive) and no-enzyme (negative) controls. All assays were performed in triplicate. IC₅₀ values (inhibitor concentration reducing PPO activity by 50%) were estimated by nonlinear regression using three-or four-parameter log-logistic models.

#### 2.3.2 Physiological Profile

Generation of the physiological profiles were performed exactly as described in Johnen et al., 2022 (27).

##### 2.3.2.1 Cell *Galium*

Freely suspended callus from *Galium mollugo* was heterotrophically cultivated in a modified Murashige-Skoog Medium (4.4g/L M&S basal medium, 30g/L sucrose, 29.7mg/L L-alanine, 3.1mg/L L-arginine-monohydrochloride, 3.8mg/L L-asparagine, 1.7mg/L L-aspartic acid, 26 mg/L gamma-aminobutyric acid, 3 mg/L L-cysteine, 0.3mg/L L-glutamine, 15.7 mg/L L-glutamic acid, 2.7mg glycine, 0.05 mg/L L-histidine-monohydrochloride, 5 mg/L L-leucine, 2 mg/L L-lysine hydrochloride, 0.05 mg/L DL-methionine, 0.05 mg/L DL-phenylalanine, 1.9 mg/L L-proline, 12.8 mg/L L-serine, 4.1 mg/L DL-threonine, 0.05 mg/L L-tyrosine, 2.3 mg/L DL-valine, 2 mg IAA, 0.1mg 2,4-D). The cells were subcultured in 7-day intervals. For screening, acetone solutions of the test compounds were pipetted into plastic tubes, and the solvent was allowed to evaporate before adding 2 mL of exponentially growing cell suspension. The tubes were shaken at 400 rpm and 25°C in the dark on a rotary shaker with an attachment for 288 test tubes. After incubation for eight days, the conductivity of the growth medium was measured using a micro-electrode (Mettler Toledo FiveEasyPlus with the electrode InoLab-752). The mean of three replicates was determined and subtracted from the value obtained before incubation. The reduction in conductivity is inversely proportional to the increase in growth. Results were expressed as percentage growth inhibition relative to untreated control.

##### 2.3.2.2 Algae

Cells of *Scenedesmus acutus* (culture collection Gottingen, 276-3a) were propagated photoautotrophically in aerated culture tubes containing an inorganic medium (1 g/L potassium nitrate, 260 mg/L disodiumhydrogenphosphat dihydrate, 740 mg/L potassium dihydrogen phosphate, 2.55 g/L magnesium sulfate heptahydrate, 25 mg/L calcium chloride hexahydrate, 0.0066 mg/L aluminum sulfate octadecahydrate, 0.099 mg/L manganese chloride tetrahydrate, 0.005 mg/L copper sulfate pentahydrate, 0.028 mg/L cobalt sulfate pentrahydrate, 0.0063 mg/L zinc sulphate heptahydrate, 0.03 mg/L boric acid, 0.002 mg/L ammonium molybdate tetrahydrate, 0.0029 mg/L Ammonium metavanadate, 0.026 mg/L

Nickel(II) sulphate hexahydrate, 0.25 mg/L potassium iodide, 0.24 mg/L potassium bromide, 0.0084 mg/L potassium bromide, 14.5 mg/L iron(III) nitrate nonahydrate, 13.3 mg/L Titriplex® III, 0.048 mg/L copper sulfate pentahydrate) and kept at 22°C under continuous light (Thorn white neon tubes, c.90 pmol m-z s-l, 400-750 nm). The bioassay was performed in plastic microtiter dishes (8.5 x 12.5 cm, NUNC) containing 24 wells. Before loading the wells with 1 mL cell suspension each, the test compounds were added in acetone, and the solvent was allowed to evaporate. The 15 additional compartments of the dishes were filled with sodium carbonate/bicarbonate buffer (0.5 ml) generating a 0.25 % partial pressure of carbon dioxide. The dishes were sealed with plastic lids and incubated on a MTS 4 shaker (IKA, Staufen, Germany) at 500 rpm under continuous light (Osram white neon tubes, c.70 pmol m-’ s-l, 400-750 nm) at 23°C. After 24 h, the contents of each well were removed, diluted and cell numbers were determined by a Coulter Counter (type ZM, Coulter Electronics, Luton, UK). Two replicates were measured, and growth inhibition was calculated as a percentage of the control.

##### 2.3.2.3 Lemna

For *L. paucicostata* cultivation and treatments, stock cultures were propagated mixotrophically in an inorganic medium containing 1 % (w/v) sucrose (KNO_3_ (400 mg/L), CaCl_2_ * 2 H_2_O (540 mg/L), MgSO_4_ * 7 H_2_O (614 mg/L), KH_2_PO_4_ (200 mg/L), Fetrilon 13 % (2,81 mg/L), MnCl_2_ * 4 H_2_O (0.415 mg/L), H_3_BO_3_ (0.5 mg/L), Na_2_MoO_4_ * 2 H_2_O (0.12 mg/L), ZnSO_4_ * 7 H_2_O (0.05 mg/L), CuSO_4_ * 5 H_2_O (0.025 mg/L) and CoCl_2_ (0.025 mg/L). The bioassay was conducted under aseptic conditions in plastic petri dishes (5 cm in diameter), which contained 15 mL medium without sucrose. The test compounds were added to the dishes in acetone solution, and the organic solvent was allowed to volatilize before four fronds were added to each dish (4 leaves per dishes). The culture dishes were then closed with plastic lids and incubated under continuous light (white LED irradiation, 70 µmol m^-2^ s^-1^) in a growth chamber at 25°C. Eight days after treatment, the increase of the area covered by the fronds in each dish was determined as the growth parameter using an image analyzing system (LemnaTec Scanalyzer; LemnaTec, Würselen, Germany). The area of fronds before incubation was subtracted from this value. Results were expressed as percentage growth inhibition relative to untreated control.

##### 2.3.2.4 Arabidopsis

Sterilized seeds (MC24 ecotype) were stratified overnight at 4°C in 48-well plates containing 250 µL half-strength Murashige & Skoog incl. Gamborgs B5 Vitamins (Duchefa: M0231.0050; 2,2 g /1 L) containing 2.5 µL of acetone (as solvent control) or 2.5 µL respective compound resulting in the following concentrations 0.001, 0.01 and 0.1 mM. 48-well plates were sealed with micropore tape and grown for 7 d in constant light at 22 °C 75% humidity. Growth inhibition and symptoms were evaluated manually.

##### 2.3.2.5 Cress germ. D and Cress germ. L

For the germination bioassay, seeds of cress (*Lepidum sativum* L) were placed in 6-well plates filled with a vermiculite substrate. Stock solutions of the test compounds in acetone were added together with 12 mL of water to reach respective test concentrations. Control seeds were moistened only with water and acetone. The plates were incubated in a growth chamber at 22 °C in the dark for 72 h. Inhibition of germination and seedling development was evaluated visually (0 = no influence, 100 = total inhibition). Afterwards, the dishes were incubated for further 3 days under light conditions (16h/8 h light:dark at 22 °C and 75% relative humidity, 230 µmol m^−2^ s^−1^ photon irradiance, 400–750 nm) and seedling development and plant symptoms were evaluated.

##### 2.3.2.6 Hill Assay

Thylakoids were isolated from shoots of 3 weeks-old plants of *Triticum aestivum* L, as follows. Plants were put 24 h in dark and subsequent steps were performed under green light. Shoots were cut and homogenized in Hill-Medium (HM) containing 50 mM tricine, 10 mM NaCl, 5 mM MgSO_4_ and 0.4 M sucrose (pH 8.0 adjusted with NaOH) using a mixer. Homogenate was filtered and centrifuged at 500 x g. Pellet was washed with suspension medium (SM) containing 50 mM tricine, 10 mM NaCl and 5 mM MgSO_4_ (pH 8.0 adjusted with NaOH) once at 500 x g and resuspended in reaction medium (RM) containing 50 mM tricine, 5 mM MgSO_4_ and 0.1 M sucrose (pH 8.0 adjusted with NaOH). For the Hill assay, isolated thylakoids (chlorophyll content 41 µg ml^−1^) in reaction medium were used. The influence of the compounds on photosynthetic electron transport in photosystem II was evaluated according to the method of Avron and Shavit (28). Briefly, the assay mixture consisted of thylakoid suspension (0.23 ml), test compound dissolved in 80% aqueous acetone + water (80 + 20 by volume; 0.05 ml), and K-ferricyanide (0.02 ml) with and without compound. During the subsequent illumination, ferrocyanide was formed in the Hill reaction if not inhibited by compound present. Then, in darkness, the ferrocyanide was allowed to react with ferric salt to form the ferrous salt which produced a complex with phenanthroline. The complex was measured photometrically by its absorption at 510 nm. Results were expressed as percent inhibition relative to untreated control and after subtraction of absorbance of the compound itself.

##### 2.3.2.7 CO_2_ assimilation

The plants of *Galium aparine* L, which had been raised under greenhouse conditions to the second-whorl stage, were cultivated in Vermiculite containing pots watered with half Linsmaier Skoog medium (KNO_3_ 950 mg/L, NH_4_NO_3_ 825 mg/L, MgSO_4_*7H_2_O 185 mg/L, CaCl_2_*2H_2_O 219 mg/L, KH_2_PO_4_ 85 mg/L, Fetrilon 13 % 46.2 mg/L, H_3_BO_3_ 3 mg/L, MnSO_4_*H_2_O 6.8 mg/L, ZnSO_4_*7H_2_O 5.3 mg/L, KJ 0.42 mg/L. Na_2_MoO_4_*2H_2_O 0.125 mg/L, CuSO_4_*5H_2_O 0.015 mg//L, CoCl_2_*6H_2_O 0.015mg/L, pH 5.8) for 5 days in a Phytotron chamber (12h day/ 12h night for 25°C / 22°C, 75% humidity). The plant treatment is a foliar spray application of a compound with the concentration of 10^-3^M. The measurement of the photosynthetic rate (µmol (CO_2_) m^-2^ s^-1^) to determine the inhibition was accomplished with a LI-COR, LI-6400XT, comparing the untreated *Galium* plant at timepoint 0h with the same plant 1 day after the compound application (24h). Additionally, epinasty and necrosis symptoms are evaluated visually. Results were expressed as percent inhibition relative to untreated control.

##### 2.3.2.8 Respiration

*Galium mollugo* were cultivated as described in Cell *Galium* (see point 1.1). 3 mL *Galium* cell suspensions were treated with compound in plastic tubes for 3-5 h in the dark on a rotary shaker. Measurement of oxygen consumption using the dissolved oxygen measuring system Mettler Toledo SevenExcellence. After 30 s of calibration the oxygen consumption (mg/L) in a time frame of 60 s was determined. The resulting consumption was used to calculate the inhibition of oxygen consumption relative to the mock treated control.

##### 2.3.2.9 Uncoupler

50 mL BY-2 suspension culture cells (7-9 days old) heterotrophically grown in Linsmaier & Skoog medium supplemented with 3 % (w/v) sucrose and 200 µg/L 2,4-D were pelleted by gravity and resuspended in 50 mL suspension buffer (50 mM HEPES, 0.5 mM CaCl2, 0.5 mM K2SO4, 10 mM Glucose, pH of 7.0 with KOH). 2 mL of cell suspension was treated with the respective compound in respective concentration in Greiner tubes and incubated for 1.5 h at 25°C in the dark while shaking (300 rpm). Cells were washed with suspension buffer.

750 µL cell suspension was incubated with 2 µg/µL JC-1 Dye in 24-well plates (Thermo Fisher, T3168) for 15 min in the dark while shaking. JC-1 exhibits potential-dependent accumulation in mitochondria, indicated by a fluorescence emission shift from green to red (29). Fluorescence emission at 590 nm at an excitation at 500 nm was measured using a plate fluorometer. Inhibition value is calculated as follows: fluorescence intensity_mock ctrl_ – fluorescence intensity_test compound_ /fluorescence intensity_mock control_ – fluorescence intensity_positive ctrl (dinoseb)_. Samples were measured in duplicates.

##### 2.3.2.10 ROS Assay

To determine the formation of reactive oxygen species (ROS), *Lemna* plants grown 7 to 9 days according to point 1.3 were treated with respective compounds and mock for 18 h and then stained with 10 μM of dihydroethidium (DHE) in 100 μM of CaCl_2_, pH 4.75, while shaking for 30 min at room temperature in the dark according to (30). DHE is a superoxide anion specific indicator that is oxidized by ROS to yield fluorescent 2-hydroxyethidium. After soaking in 100 μM of CaCl_2_ for 5 min to remove the residual dye, fluorescence of 2-hydroxyethidium in *Lemna* roots was observed using an Olympus BX61 epifluorescence microscope (Hamburg, Germany). ROS formation was evaluated visually (0 = no formation, 100 = strong formation).

##### 2.3.2.11 Chlorophyll fluo

For *L. paucicostata* cultivation and treatments, stock cultures were propagated mixotrophically in an inorganic medium containing 1 % (w/v) sucrose (KNO_3_ (400 mg/L), CaCl_2_ * 2 H_2_O (540 mg/L), MgSO_4_ * 7 H_2_O (614 mg/L), KH_2_PO_4_ (200 mg/L), Fetrilon 13 % (2,81 mg/L), MnCl_2_ * 4 H_2_O (0.415 mg/L), H_3_BO_3_ (0.5 mg/L), Na_2_MoO_4_ * 2 H_2_O (0.12 mg/L), ZnSO_4_ * 7 H_2_O (0.05 mg/L), CuSO_4_ * 5 H_2_O (0.025 mg/L) and CoCl_2_ (0.025 mg/L)). The bioassay was conducted under aseptic conditions, using 15 mL medium without sucrose. The bioassay was performed in plastic microtiter dishes (8.5 x 12.5 cm, NUNC) with 24 wells. Loading the wells with 1 mL cell suspension and approx. 12 fronds of *Lemna*. Effects on chlorophyll fluorescence (fluorescence quantum yield, Y(II)) were measured after 24 h compound treatment using an Imaging Pulse-Amplitude-Modulation (PAM) M-Series system (Walz, Effeltrich, Germany). Measurements were taken after 5 minutes of darkness with the following settings: Meas.light=1, act. Light=2. Results were expressed as percent inhibition relative to untreated control.

##### 2.3.2.12 ATP content

Growth and treatment were performed according to Chlorophyll fluorescence (point 1.11). ATP content was measured with the ATP Determination Kit (A22066, Thermo Fisher Scientific,Grand Island, NY14072) according to manufacturer’s instructions using 9-12 mg *L. paucicostata* extract in tricine buffer. Results were expressed as percentage growth inhibition relative to untreated control and fresh weight.

##### 2.3.2.13 Neural Network Modeling of Physiological Profiling Data

A variational autoencoder was fitted to the results of 14 types of physiological assays for herbicide compounds which were assessed at different concentrations as described in Johnen et al., 2022 (10.1002/ps.7004). Concentrations were treated as different features, resulting in a total of 34 features. Samples per compound/assay/concentration combinations were aggregated by median. The autoencoder was semi-supervised with a classification network attached to the latent samples and learned a latent space with eight dimensions.

Multilabel classification was applied to allow for non-mutually exclusive MoA predictions.

Eight classes of modes of action, along with an’Unknown Mode of Action’ class were used as targets. Monte-Carlo dropout was applied in the classification network. The uncertainty resulting from the samples in latent space along with the Monte-Carlo dropout samples of the classification network allowed for assessments of uncertainty of predicted MoA labels. Reference sample contributions to the cross entropies MoA class were inversely weighted by the number of positive examples over negative examples, for their respective MoA classes. Missing values for input features and labels were imputed with zeros but were masked in the respective loss function terms. The Kullback-Leibler Divergence (KLD) and classification loss terms were balanced using weight parameters in the loss function. The KLD and classification weights were slowly ramped up during training, to help the learning. The test compound fendioxypyracil was part of the test set of unknown mode-of-actions.

### 2.4 Herbicidal Test Methods

#### 2.4.1 POST- Emergence Greenhouse Trials

The active ingredients for the POST-Emergence greenhouse trials were selected from the most used PPO inhibitors (HRAC E, 14) in US and Brazil Soy fields: saflufenacil (N-Phenyl-imide, manufacturer: BASF), trifludimoxazin (N-Phenyl-imide, BASF), tiafenacil (N-Phenyl-imide, Nufarm), Flumioxazin (N-Phenyl-imide, Sumitomo).

The key broadleaf weed species and the grass species that were tested in the greenhouse trials are described in Table 1, including the EPPO Codes (former Bayer Codes, European and Mediterranean Plant Protection Organization). All used seeds are an own production in BASF’s Germany site at Limburgerhof. Application was done at BBCH 12/13.

**Table 1:** Investigated monocotic weeds and dicot weeds of the Post-Emergence Trial.

| **EPPO Code** | **Scientific name** | **English name** | **Authority** |
| --- | --- | --- | --- |
| ABUTH | *Abutilon theophrasti* | velvet leaf | Medicus |
| AMAPA | *Amaranthus palmeri* | palmer amaranth | Watson |
| AMARE | *Amaranthus retroflexus* | redroot pigweed | Linnaeus |
| AMATA | *Amaranthus x tamariscinus* | tall amaranth | Nuttall |
| AMBEL | *Ambrosia artemisiifolia* | common ragweed | Linnaeus |
| BIDPI | *Bidens pilosa* | hairy beggarticks | Linnaeus |
| CAPBP | *Capsella bursa-pastoris* | shepherd's purse | (Linnaeus) Medicus |
| CHEAL | *Chenopodium album* | common lambsquarters | Linnaeus |
| COMBE | *Commelina benghalensis* | Bengal day flower | Linnaeus |
| EPHHL | *Euphorbia heterophylla* | wild poinsettia | Linnaeus |
| ERICA | *Erigeron canadensis* | Canadian horseweed | Linnaeus |
| GALAP | *Galium aparine* | cleavers | Linnaeus |
| IPOHE | *Ipomoea hederacea* | morning glory | Jacquin |
| KCHSC | *Kochia scoparia* | kochia | (Linnaeus) Schrader |
| MATCH | *Matricaria chamomilla* | wild chamomile | Linnaeus |
| PAPRH | *Papaver rhoeas* | common poppy | Linnaeus |
| RAPRA | *Raphanus raphanistrum* | wild radish | Linnaeus |
| SEBEX | *Sesbania herbacea* | coffeebean | (Miller) McVaugh |
| SIDRH | *Sida rhombifolia* | common sida | Linnaeus |
| SOLNI | *Solanum nigrum* | black nightshade | Linnaeus |
| XANST | *Xanthium strumarium* | common cocklebur | Linnaeus |
| ALOMY | *Alopecurus myosuroides* | blackgrass | Hudson |
| AVEFA | *Avena fatua* | wild oat | Linnaeus |
| BRARU | *Brachiaria decumbens* | Surinam grass | Stapf |
| BROST | *Bromus sterilis* | poverty brome | Linnaeus |
| CYPIR | *Cyperus iria* | rice flatsedge | Linnaeus |
| DIGSA | *Digitaria sanguinalis* | crabgrass | (Linnaeus) Scopoli |
| ECHCG | *Echinochloa crus-galli* | barnyard grass | (Linnaeus) Palisot de<br>Beauvois |
| ECHCO | *Echinochloa colonum* | small barnyard grass | (Linnaeus) Link |
| ELEIN | *Eleusine indica* | goosegrass | (Linnaeus) Gärtner |
| ERBVI | *Eriochloa villosa* | woolly cupgrass | (Thunberg) Kunth |
| LEFFA | *Leptochloa fusca* subsp.<br>*fascicularis* | bearded sprangletop | (Lamarck) Snow |
| LOLMU | *Lolium multiflorum* | Italian ryegrass | Lamarck |
| LOLPE | *Lolium perenne* | English ryegrass | Linnaeus |
| PANDI | *Panicum dichotomiflorum* | fall panicum | Michaux |
| SETFA | *Setaria faberi* | giant foxtail | Herrmann |
| SETIT | *Setaria italica* | foxtail millet | (Linnaeus) Palisot de<br>Beauvois |
| SETVE | *Setaria verticillate* | bristly foxtail | (Linnaeus) Palisot de<br>Beauvois |
| SETVI | *Setaria viridis* | green foxtail | (Linnaeus) Palisot de<br>Beauvois |
| SORHA | *Sorghum halepense* | Johnson grass | (Linnaeus) Persoon |

\* from EPPO Global Database: https://gd.eppo.int/

The plants were cultivated using standard methods in Limburgerhof soil (slightly loamy sand soil, clay 6,9% dm; loam 16,6% dm; sand 76,5% dm, organic matter (OM) 1,38% dm; pH 7,4). The plant pots had a diameter of 9 cm at the broadest point and contained approximately 313 cm^3^ of soil (standard pot size). Monocot weeds were cultivated directly in the plant pots. Dicot weeds were cultivated in propagation soil (pH 5,6; N 14%, P_2_O_5_ 16%, K_2_O 18%, Fe 0,09%) and transplanted into pots with Limburgerhof soil after emergence.

The plants were treated with specific formulated active ingredients at various application rates to evaluate their responses to different dosages. The application was carried out under controlled conditions to facilitate a clear distinction between the active compounds and to manage the various weed species effectively. An initial trial aimed to establish suitable application rates. Given that most of the compounds are UV-dependent, significantly lower rates were employed in greenhouse trials compared to field rates. For consistency, all PPO inhibitors were applied at a uniform rate (Table 2).

**Table 2.** Application conditions for active ingredients in the Post-Emergence Trial. All actives were applied with 1% MSO (Lecitech) as adjuvant and a water volume of 200 L/ha.

| Active Ingredient | Formulation* | Rate (g ai ha-1) |  |  |  |
| --- | --- | --- | --- | --- | --- |
| Water | Control |  |  |  |  |
| Saflufenacil | 342 g L-1 SC | 8 | 4 | 2 | 1 |
| Trifludimoxazin | 500 g L-1 SC | 8 | 4 | 2 | 1 |
| Flumioxazin | 51% WG | 8 | 4 | 2 | 1 |
| Tiafenacil | 50 g L-1 ME | 8 | 4 | 2 | 1 |
| Fendioxypyracil | 342 g L-1 SC | 8 | 4 | 2 | 1 |

\*SC: Suspension concentrate, WG: Water dispersible granules, ME: Microemulsion, SL: Soluble liquid concentrate, g ai ha-1: gram active ingredient per hectare

The POST-Emergence trial was repeated 4 times in terms of having enough replication for each evaluation of each rate. The application volume was standardized at 200 liters per hectare, with 0.5% methylated seed oil (MSO) used as an adjuvant. All applications were conducted using a flat spray nozzle from the XR Teejet 110015VS series. After treatment, the solvents and water were allowed to evaporate from the plants for 30 minutes in a separate tunnel with an airflow of 3000 m-3 h. Subsequently, the plants were transferred to greenhouses tailored to the required growing conditions. The trials utilized three different greenhouses: a warm house (22-24°C, mean humidity 57%), a cold house (18-21°C, mean humidity 64%), and a cold cabin (12-14°C, mean humidity 83%). Each greenhouse was illuminated with photosynthetically active radiation (PAR; 380 – 780 nm) from 10:00 p.m. to 4:00 a.m., in addition to natural daylight.

Irrigation for the plants was conducted using specially prepared water that included nutrients tailored to their growth stage, biomass availability, and water needs. The irrigation water was prepared by diluting 1 per mile of the liquid fertilizer “Kamasol brilliant Grün 10-4-7®” in tap water.

Plant damage was assessed at 7- and 20-days post-application of the active ingredients. The evaluation involved a visual inspection of the above-ground parts of the plants, with damage quantified as a percentage of Plant Damage Compared to Untreated Control (PDCU) using a scale ranging from 0 to 100, including increments of 2 (0%, 5%, 10%, 15%,…, 90%, 95%, 98%, 100%). A PDCU value of 0% indicated no damage, while 100% indicated complete plant death. For the analysis of the rating data collected, the statistical software R was utilized. The analysis of variance (ANOVA) technique, as outlined by Stahle and Wold in 1989, was employed to identify differences in means. When significant differences were noted in the ANOVA results, the means were categorized into distinct groups following the method described by Scott and Knott, using a significance level (α) of 0.05. The clustering analysis method developed by A. Scott and M. Knott (31) was applied to group the variants into cohesive and homogeneous categories.

## 3. Results and discussion

### 3.1 Synthesis

Fendioxypyracil can be synthesized starting from 3-chloro-2,5,6-trifluoropyridine (Scheme 2). In a sequential one-pot nucleophilic substitution reaction using sodium azide followed by 2-(benzyloxy)phenol, the respective aryl-ether azide is formed. Reduction of the azide to the amine using zinc and subsequent reaction with ethyl chloroformate generates the respective carbamate building block **8**. From the carbamate the uracil ring is constructed by employing ethyl 3-amino-trifluorobutenoate followed by N-methylation. Deprotection of the benzyl-group and alkylation with ethyl bromoacetate provides then fendioxypyracil (**5**) with very good yields.

### 3.2 Physiological profiling of fendioxypyracil indicates inhibition of PPOs as MoA

Physiological profiling for the identification of herbicidal modes of action has been successfully used and refined for decades (30, (27, 32). The MoA identification is based on the generation of an inhibition fingerprint from an array of different assays that also includes the assessment of the compound-induced symptoms. To identify its mode of action, we generated a physiological profile (P-Profile) for fendioxypyracil (Fig 2A). Fendioxypyracil induced rapid necrosis and led to substantial inhibition in plant tissues that were incubated in light, as observed in *Lemna paucicostata*, algae and cress seedlings. In addition, the germination and growth of light incubated *Arabidopsis thaliana* seedlings was strongly inhibited by fendioxypyracil. In contrast, heterotrophically, dark grown *Galium* cell suspensions were only affected at high concentrations and cress germination in darkness was only mildly inhibited. Carbon dioxide (CO_2_) assimilation in *Galium* plants was strongly inhibited by fendioxypyracil. In contrast, fendioxypyracil only slightly affected the Hill reaction and the chlorophyll fluorescence assay, which uses the quantum yield (Y(II)), as an indicator for how much absorbed light energy is used for photochemical reactions in photosystem II (PSII). The above-described results show that even though the overall photosynthesis is affected by fendioxypyracil, it is not based on the direct inhibition of the PSII but it is light dependent. No or only mild effects were observed in the respiration assay as measured through oxygen consumption in heterotrophic *Galium* cell suspensions and on mitochondrial membrane potentials in *Lemna* roots. Fendioxypyracil treatment led to rapid necrosis, chlorosis and inhibition of root growth in *Lemna*; necrosis and inhibition of root growth in the cress germination assay and in *Galium* plants used for the analysis of CO_2_ accumulation.

**Figure 2:**
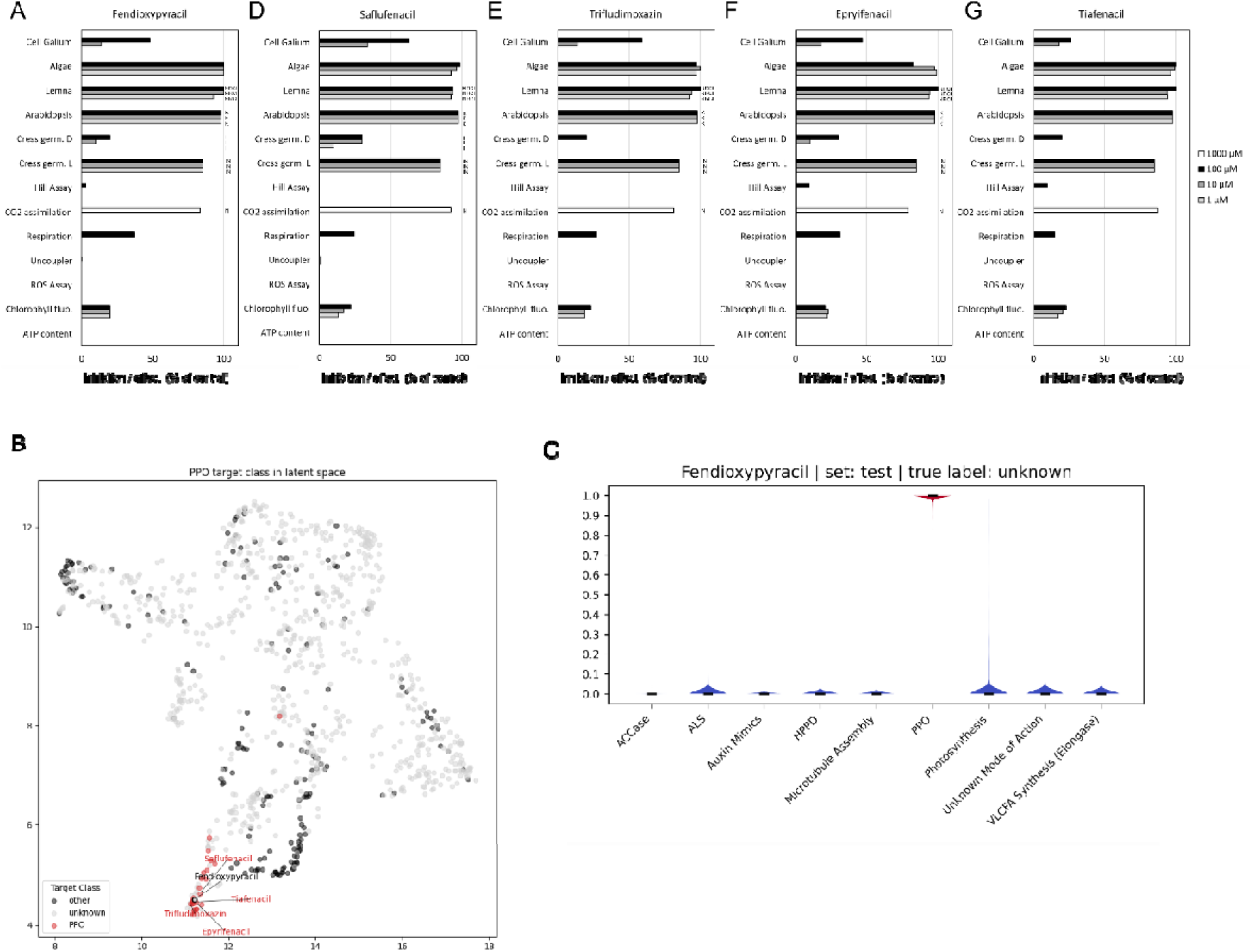
(A, D-G) **(A)** Physiological profiles of fendioxypyracil, saflufenacil, trifludimoxazin, epyrifenacil and tiafenacil. Effect of respective compounds on heterotrophic *Gallium mollugo* cell suspensions, green algae *Scenedesmus obliquus*, *Lemna paucicostata* plants, *Arabidopsis thaliana* seedling morphology, germination of cress (*Lepidium sativum* L.) under dark/light conditions, the Hill reaction in isolated *Triticum aestivum* chloroplasts, CO2 assimilation in *G. mollugo* plants, oxygen consumption (respiration) in heterotrophic *G. mollugo* cell suspensions, uncoupler activity and accumulation of reactive oxygen species (ROS) in L. *paucicostata* root tissue, chlorophyll fluorescence, and ATP content are shown. Uncoupler activity and ROS accumulation are shown as effect compared with control, while values of all other assays are expressed as percentage of inhibition. Letters indicate physiological symptoms observed: rapid necrosis (NR), necrosis (N), root growth inhibition (I), chlorosis (C) and inhibition of germination (K) **(B)** Latent Space learned by a variational autoencoder model using P-Profile results generated for a diversity of herbicides. Dimensional reduction of the latent space has been performed by UMAP. Pesticides with unknown target class are shown in light grey along pesticides with known target classes, highlighted in red for known PPO (protoporphyrinogen oxidase) inhibitors, namely saflufenacil, trifludimoxazin, epyrifenacil and tiafenacil, and in dark grey for eight additional reference target classes detailed in C. The test compound fendioxypyracil is displayed as light grey dot with a black border. Fendioxypyracil aligns with known PPO inhibitors in the latent space **(C)** Calculated probabilities (y axis) computed by a multilabel classification network including nine target classes (x axis). Prediction uncertainty has been assessed by Monte Carlo dropout and is shown as spread (y axis).

**Figure 3.**
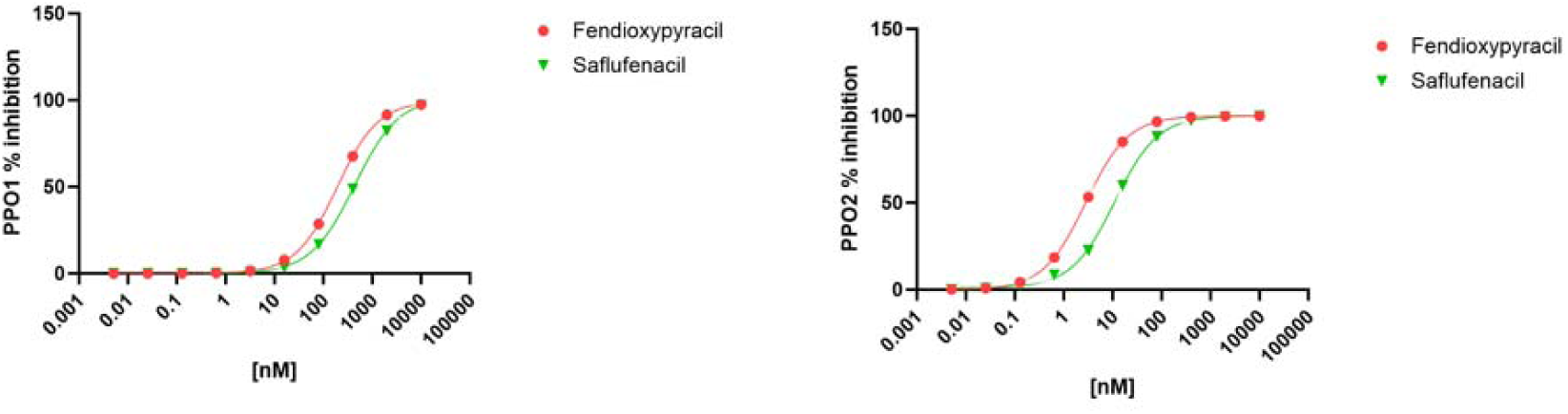
Dose–response curves showing percent inhibition of (left) PPO1 and (right) PPO2 by fendioxypyracil (red circles) and Saflufenacil (green triangles). Enzymatic activity was measured across a range of inhibitor concentrations (0.001–100,000 nM), and percent inhibition was calculated relative to the activity of enzyme with no herbicide (positive control). Data points represent mean values from three replications, and curves were fitted using a four-parameter logistic model to estimate IC₅₀ values (Table 4).

To identify the mode of action, we analyzed the P-Profile result by application of a semi-supervised variational autoencoder model. The effects of the test compound fendioxypyracil were investigated first by exploration of the learned latent space and second by calculated probabilities for nine target classes (eight known modes of action and ‘Unknown Mode of Action’) in the multilabel classification prediction. The latent space which has been learned over results derived from a diversity of herbicides including commercially-relevant reference herbicides listed and classified by the herbicide resistance action committee (based on herbicides listed on the HRAC poster 2024 (33), is shown in Figure 2B (latent space). The eight dimensions of the learned latent space have been reduced to two dimensions by UMAP (Uniform Manifold Approximation and Projection) for visualization purpose.

Fendioxypyracil (treated as test compound in the model prediction) aligns within a cluster of established PPO inhibitors, saflufenacil, trifludimoxazin, epyrifenacil and tiafenacil (Figure 2B, latent space) (4). The calculated probabilities for eight different known MoA classes and ‘Unknown Mode of Action’ modeled are shown in Fig. 2C (probabilities) and in Table 3 and indicate a probability of over 99% for the MoA PPO inhibition and a very low probability of all other target classes. The very low uncertainty of the prediction as modeled by Monte Carlo dropout and displayed by the spread of the probabilities in Fig. 2C (probabilities) indicates a high reliability of the prediction results. Both the location of fendioxypyracil within the latent space (Figure 2B, latent space) as well as the calculated probabilities of the classification model (Fig. 2C, Table 3, probabilities) indicated PPO inhibition as MoA.

**Table 3.**
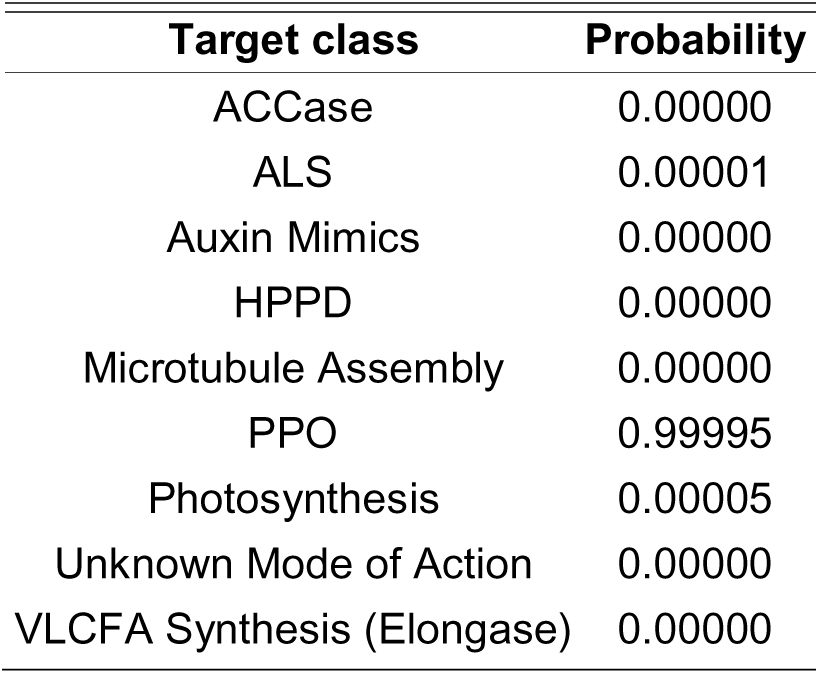
Calculated probabilities computed by a multilabel classification network including nine target classes reveal PPO as the most likely MoA of fendioxypyracil and a very low probability for other modes-of-action target classes.

For the known PPO inhibitor saflufenacil a P-Profile was already reported by Grossmann et al., 2010 (4) albeit in a slightly refined setup as exploited here. Nevertheless, the inhibitory effects and symptoms in the P-Profile of saflufenacil (i.e. light dependent inhibitory effects on photosynthesis without affecting the PSII and induction of necrotic symptoms, https://doi.org/10.1614/WS-D-09-00004.1) are resembling the profile of fendioxypyracil.

Furthermore, we generated profiles with saflufenacil, trifludimoxazin, epyrifenacil and tiafenacil identified in the PPO cluster (Fig. 2B) in the current P-Profile setup and to compare the profiles to that induced by fendioxypyracil. The inhibition profile and the observed symptomology induced by fendioxypyracil were highly similar to those of saflufenacil, trifludimoxazin, epyrifenacil and tiafenacil (Fig 2A, D-G) corroborating the outcome of the classification model. Collectively, these analyses indicated that the MoA of fendioxypyracil is based on the inhibition of PPOs *in planta*.

### 4.2 Fendioxypyracil inhibits both the *Amaranth tuberculatus* PPO1 and PPO2 wild type enzymes *in vitro*

Enzyme inhibition assays confirmed that both PPO1 and PPO2 from *A. tuberculatus* are strongly inhibited by fendioxypyracil and the benchmark compound saflufenacil, as well-characterized PPO inhibitor (34) (Table 4). For PPO1, fendioxypyracil displayed an IC₅₀ of 195 nM, whereas saflufenacil was less potent, with an IC₅₀ of 412 nM. This indicates that fendioxypyracil requires roughly half the concentration of saflufenacil to achieve comparable inhibition of PPO1 activity. A more pronounced difference was observed with PPO2, where fendioxypyracil exhibited an IC₅₀ of only 2.8 nM compared with 11 nM for saflufenacil. Thus, fendioxypyracil inhibited PPO2 with approximately four-fold greater potency than the benchmark herbicide. At a saturating concentration of 10 µM, both compounds fully inhibited PPO1 and PPO2 activity, resulting in 100% loss of enzymatic activity. Taken together, these results demonstrate that fendioxypyracil is a highly effective inhibitor of both PPO isoforms *in vitro*. While saflufenacil confirmed its expected inhibitory activity, fendioxypyracil consistently showed stronger potency, particularly against PPO2, highlighting its potential as a powerful PPO-targeting herbicide.

**Table 4.** *In vitro* inhibition of *Amaranthus tuberculatus* PPO1 and PPO2 wild-type enzymes by fendioxypyracil and the benchmark herbicide saflufenacil. IC₅₀ values (nM) represent the herbicide concentration required to reduce enzyme activity by 50%, as estimated from nonlinear regression of dose–response curves. Saflufenacil was included as a reference compound. % inhibition values were determined at a saturating concentration of 10 µM, at which both enzymes were fully inhibited. Each value is an average of three replications (P < 0.0001).

| PPO isoform | Enzyme backbone | Fendioxypyracil IC50[nM]<br>P<0.0001 | Saflufenacil IC50[nM]<br>P<0.0001 | Fendioxypyracil % Inhibition at saturation | Saflufenacil % Inhibition at saturation |
| --- | --- | --- | --- | --- | --- |
| PPO1 | AMATU | 195 | 412 | 100 | 100 |
| PPO2 | AMATU | 2.8 | 11 | 100 | 100 |

### 4.3 Weed performance of Fendioxypyracil in Post-emergence application in the greenhouse and confirmation in field trials

The post-emergence greenhouse trials demonstrated that fendioxypyracil is highly effective against a broad spectrum of weed species, including both grasses and broadleaf weeds. At a rate of 16 g active ingredient per hectare, fendioxypyracil controlled more than 80 weed species, as shown in Table 5 (fig 4 and 5). This broad activity was confirmed in multiple field trials, highlighting its robust performance under practical conditions (fig 7).

**Figure 4:**
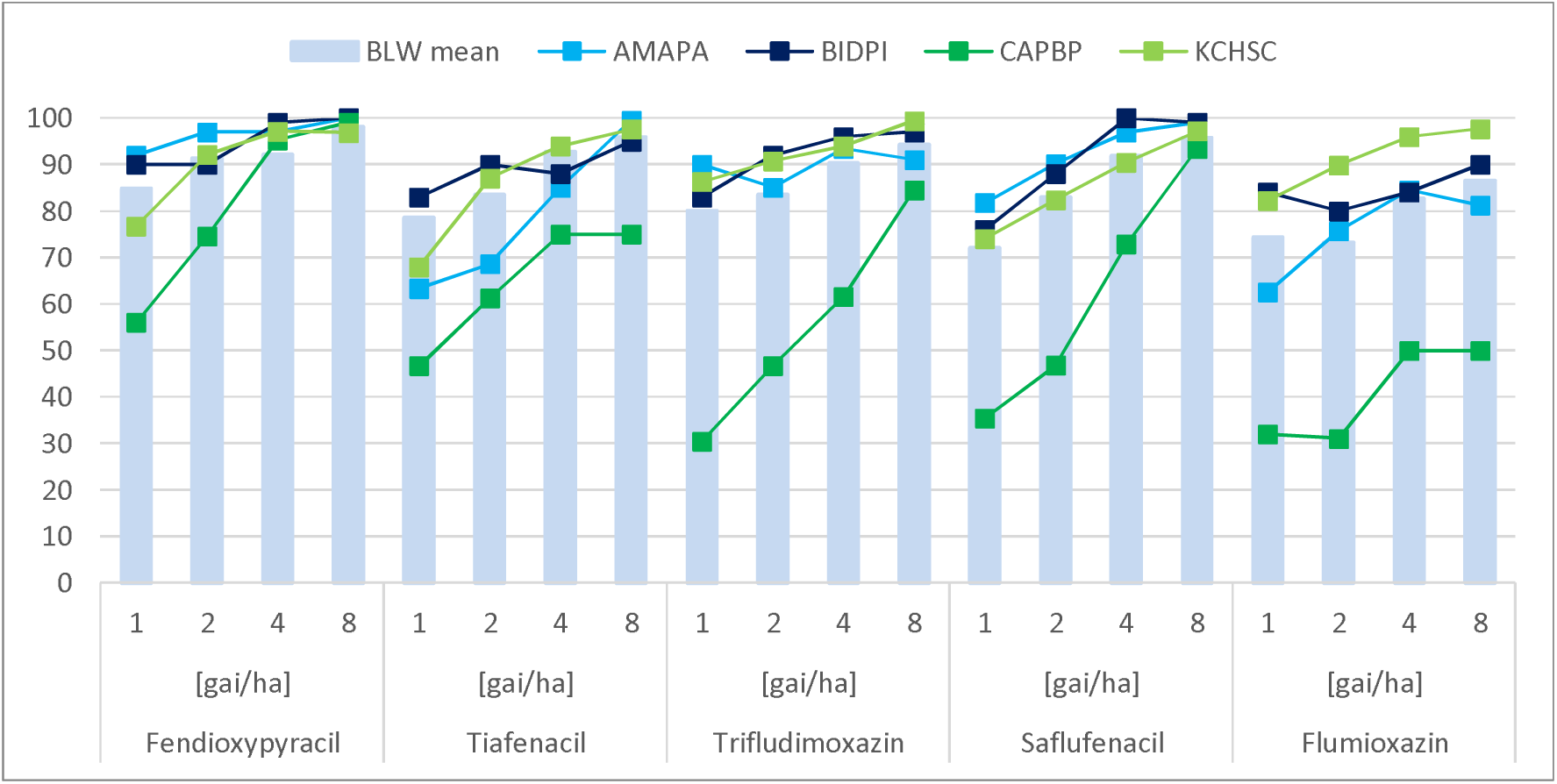
Broadleaf weed (BLW) control in greenhouse. Represented is the BLW average control as well as activity against specific BLW, such as AMAPA, BIDPI, CAPBP, KCHSC. The assessment was done 20 days after treatment

**Figure 5:**
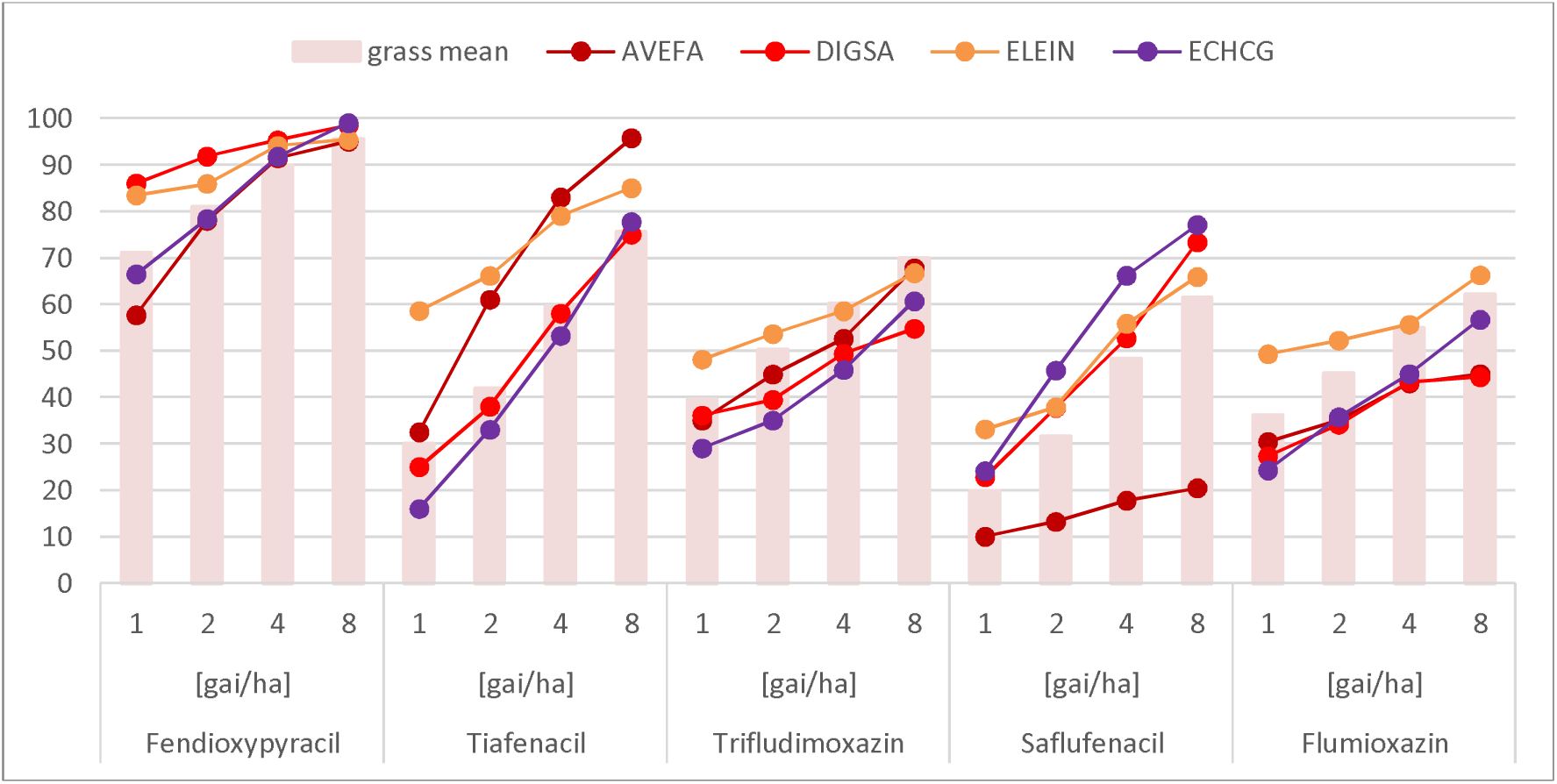
Grass control after greenhouse application. Represented is the mean grass control as well as activity against AVEFA as cold season grass, DIGSA, ELEIN and ECHCG as warm season grasses. The assessment was done 20 days after treatment.

**Figure 6:**
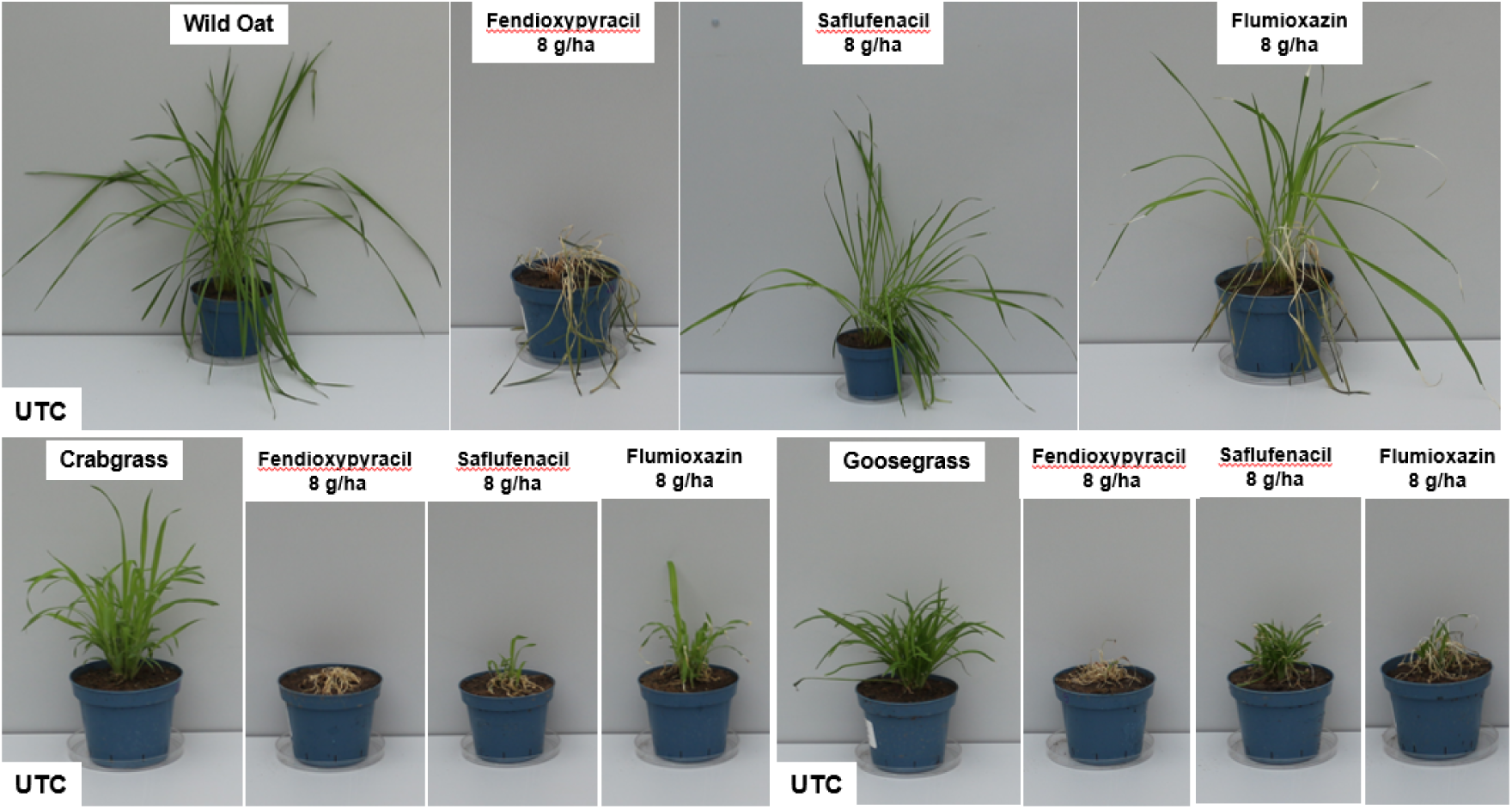
Grass activity in glasshouse pot trial. Post-emergence application on wild oat, crabgrass and goosegrass. Comparison of fendioxypyracil, saflufenacil and flumioxazin. Spray volume 200 L/ha with 0.5 % MSO, 20 Days after treatment

**Figure 7:**
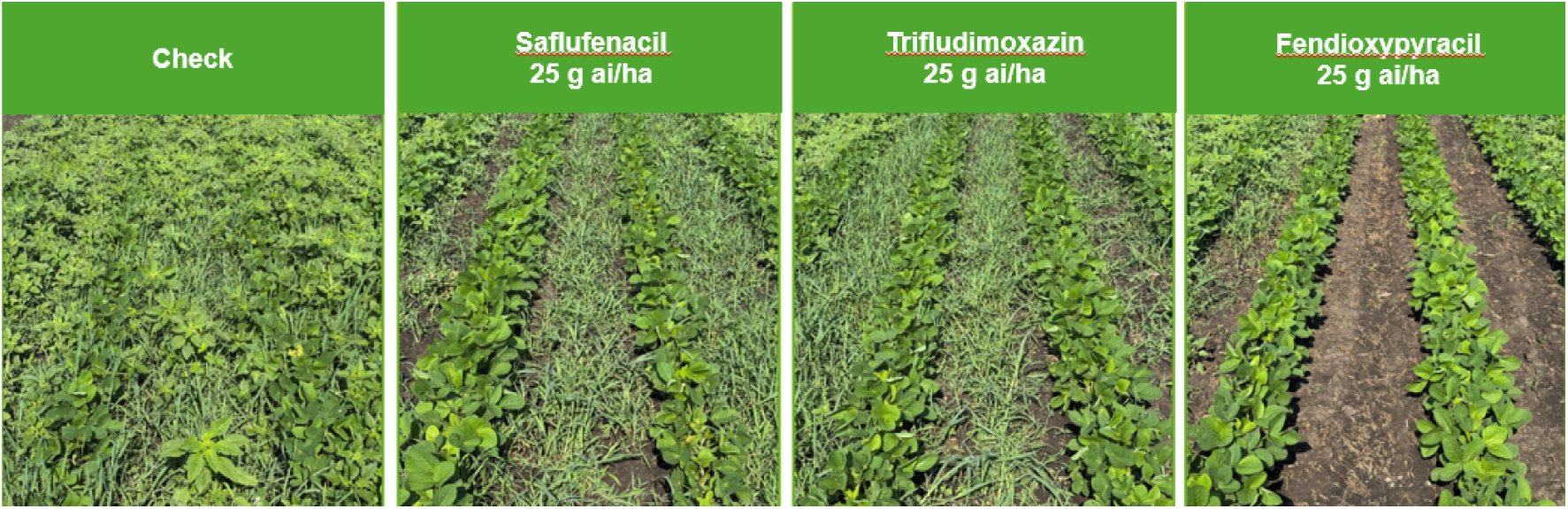
Field Trials in Seymour, IL, USA. Pre-Plant Burndown application, using 200 l/ha with 0,5 % MSO and 1 % AMS. 9 days after application. Control of AMASS, ECHCG and SETSS

**Table 5.** Examples of weeds controlled by fendioxypyracil in Greenhouse at 16 g/ha.

| Weed Type |  |
| --- | --- |
| **Broadleaf weeds** | Abutilon theophrasti, Amaranthus palmeri, Amaranthus retroflexus, Amaranthus x tamariscinus, Ambrosia artemisiifolia, Bidens pilosa, Capsella bursa-pastoris, Centaurea cyanus, Chenopodium album, Cirsium arvense, Commelina benghalensis, Convolvulus arvensis, Euphorbia heterophylla, Eriochloa villosa, Erigeron canadensis, Galium aparine, Geranium dissectum, Geranium pusillum, Ipomoea hederacea, Ipomoea lacunose, Kochia scoparia, Lamium amplexicaule, Lamium purpureum, Matricaria chamomilla, Matricaria inodora, Oryza rufipogon, Panicum dichotomiflorum, Panicum miliaceum, Papaver rhoeas, Physalis ixocarpa, Polygonum convolvulus, Portulaca oleracea, Portulaca oleracea subsp. sativa, Raphanus raphanistrum, Salsola kali subsp. ruthenica, Sesbania herbacea, Sida rhombifolia, Sinapis alba, Sinapis arvensis, Solanum nigrum, Sonchus oleraceus, Stellaria media, Taraxacum officinale, Thlaspi arvense, Veronica persica, Viola arvensis, Xanthium strumarium |
| **Grasses** | Agrostis stolonifera, Alopecurus myosuroides, Apera spica-venti, Avena fatua, Brachiaria decumbens, Brachiaria plantaginea, Bromus inermis, Bromus sterilis, Cyperus esculentus, Cyperus iria, Digitaria sanguinalis, Echinochloa crus-galli, Echinochloa colonum, Eleusine indica, Eriochloa villosa, Ischaemum rugosum, Leptochloa chinensis, Leptochloa fusca subsp. fascicularis, Leptochloa mucronate, Lolium multiflorum, Lolium perenne, Oryza rufipogon, Panicum dichotomiflorum, Panicum miliaceum, Paspalum notatum, Phalaris canariensis, Poa annua, Rottboellia cochinchinensis, Setaria faberi, Setaria italica, Setaria verticillate, Setaria viridis, Sorghum halepense |

A key finding is fendioxypyracil’s exceptional control of grass weeds—such as wild oat, barnyard grass, crabgrass, and goosegrass—by foliar application in greenhouse pot tests (fig 5 and 6). This level of grass control is unique among PPO inhibitors, which typically show limited efficacy against grasses. In comparative trials in the greenhouse, fendioxypyracil was tested alongside other PPO-inhibiting herbicides (saflufenacil, trifludimoxazin, flumioxazin, tiafenacil), all applied at identical rates and with the same adjuvant. The results showed that all PPO inhibitors provided strong control of broadleaf weeds, with minimal differences in efficacy (Figure 4). However, fendioxypyracil stood out for its consistently high efficacy, even at the lowest tested dose of 1 g ai/ha on the greenhouse, and for its flat dose-response curve, indicating reliable performance across a range of application rates. Fendioxypyracil achieved high average control of broadleaf weeds, matching or exceeding the performance of other PPO inhibitors. Specific weeds such as *Amaranthus palmeri* (AMAPA), *Bidens pilosa* (BIDPI), *Capsella bursa-pastoris* (CAPBP), and *Kochia scoparia* (KCHSC) were effectively controlled (fig 4).

Fendioxypyracil provided superior control of both cold-season grasses (e.g., *Avena fatua*, AVEFA) and warm-season grasses (e.g., *Digitaria sanguinalis*, DIGSA; *Eleusine indica*, ELEIN) compared to other PPO inhibitors. Notably, only tiafenacil approached similar levels of grass control, but fendioxypyracil maintained a steadier response rate and higher overall efficacy (Figure 5).

Fendioxypyracil has demonstrated outstanding efficacy in global field trials. Field data showed (fig 7) that fendioxypyracil achieves rapid and thorough weed control at low use rates (25 g ai/ha) in pre-plant burndown applications, outperforming other PPO inhibitors, especially in grass weed management (ECHCG and *Setaria species*, SETSS). In the US, fendioxypyracil delivered near-complete control of AMAPA a major driver of herbicide resistance issues. These results position fendioxypyracil as a valuable tool for sustainable weed management in major cropping systems, offering growers a new solution for pre-plant burndown and resistance management strategies.

In summary, fendioxypyracil offers outstanding post-emergence weed control, particularly for grass species a significant advancement for PPO-inhibiting herbicides. Its broad-spectrum activity, reliable efficacy at low dose rates, and unique strength against grasses makes it a valuable tool for sustainable weed management in major crops.

## 4. Conclusion

Fendioxypyracil is a novel PPO-inhibiting foliar herbicide with strong PPO target inhibition, offering broad spectrum weed control (grass and broadleaf weeds) at low use rates. These key attributes will make fendioxypyracil a useful (effective?) preplant burndown herbicide for broad acre crops such as soybeans and corn. Future PPO inhibitor tolerant crops will enable over-the-top applications of fendioxypyracil, thereby broadening its use in future weed management programs. Fendioxypyracil is being developed in many key agricultural countries such as USA, Brazil and Argentina. It is expected to be commercially available in the first countries around the end-2020s, contributing to global food production in the midterm future. Resistance testing is in progress and aims to assess fendioxypyracil activity against PPO-resistant weed populations target site mutant PPO enzymes, and a comprehensive evaluation of resistance mechanisms and management implications will be the objective of a subsequent study.

## 5. Acknowledgements

We would like to thank Susanne Knauer, Misha Manuchehri Byrd, Simone Huber, Sarina Rühm und Stefanie Zimmermann for the experimental support. This research did not receive any specific grant from funding agencies in the public, commercial, or not-for-profit sectors.

## Schemes and Figures

**Scheme 1:**
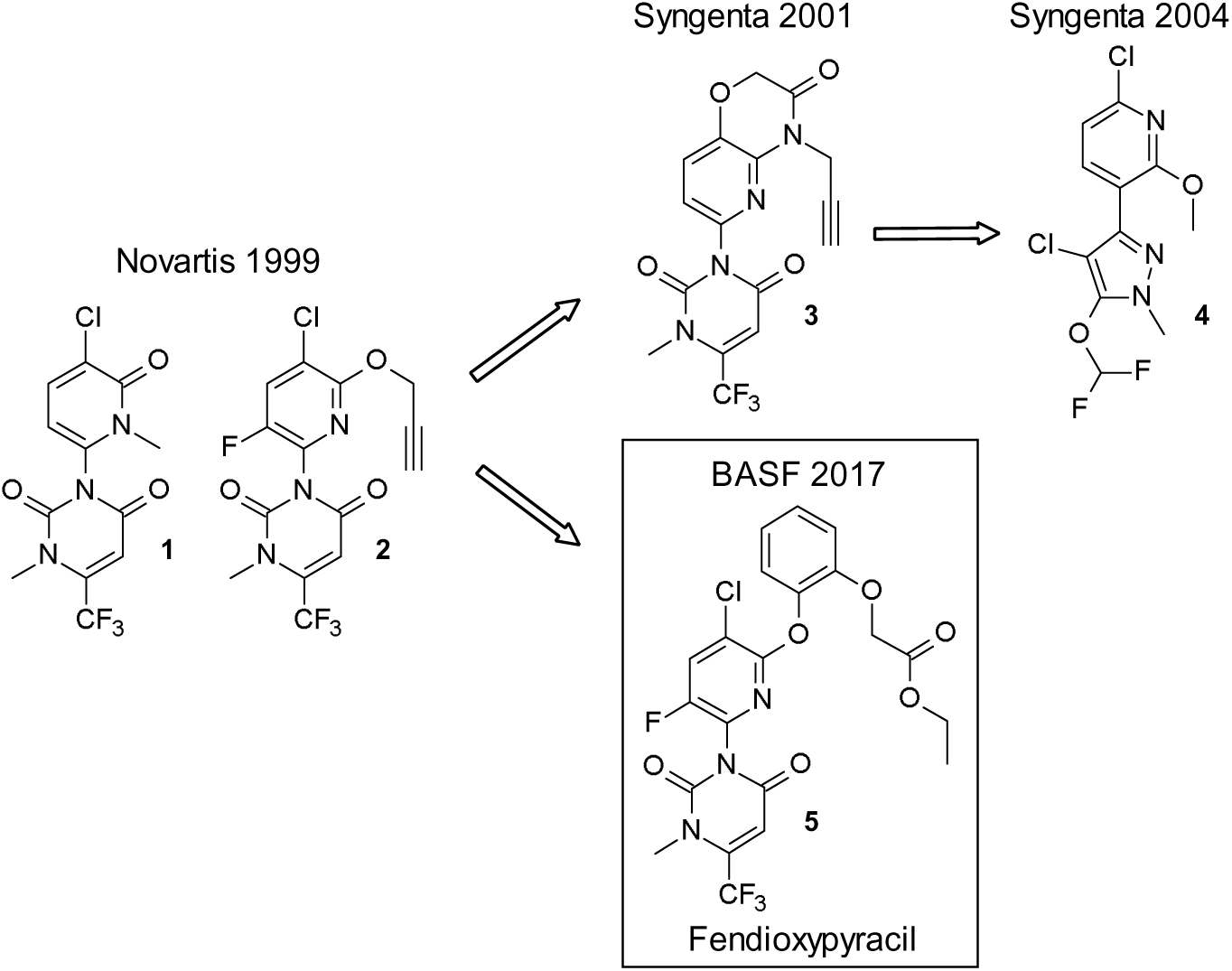
Discovery of PPO herbicide structures with a central pyridine core

**Scheme 2:**
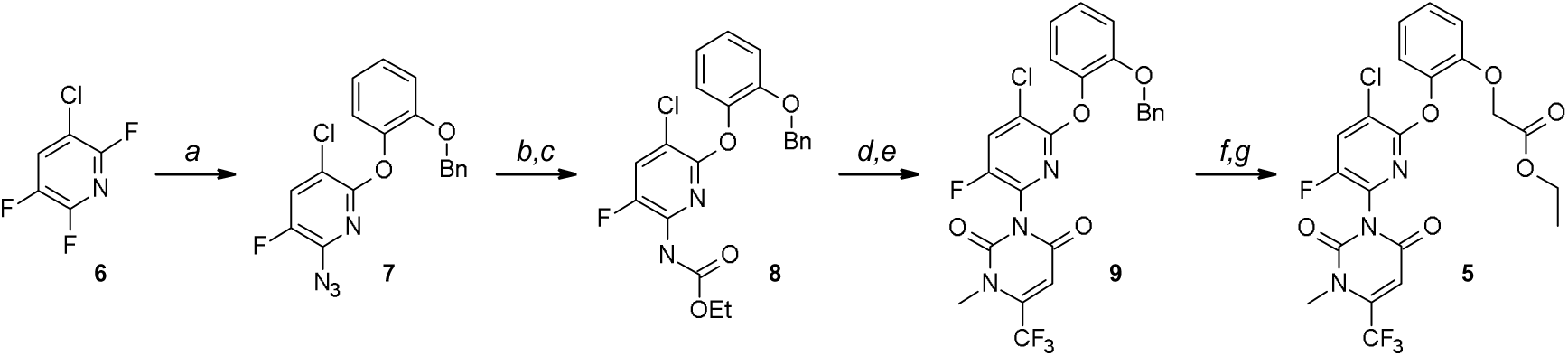
Fendioxypyracil synthesis: (a) i) NaN3, ii) 2-(Benzyloxy)phenol; (b) Zn, NH4Cl; (c) ethyl chloroformate; (d) ethyl (E)-3-amino-4,4,4-trifluoro-but-2-enoate; (e) MeI; (f) AlCl3; (g) ethyl bromoacetate.

